# CRISPR/Cas9 gene editing to generate *Drosophila* LexA lines in secondary school classes

**DOI:** 10.1101/2023.06.22.546134

**Authors:** Anne E. Rankin, Elizabeth Fox, Townley Chisholm, Nicole Lantz, Arjun Rajan, William Phillips, Elizabeth Griffin, Jaekeb Harper, Christopher Suhr, Max Tan, Jason Wang, Alana Yang, Ella S. Kim, Naa Kwama A. Ankrah, Praachi Chakraborty, Alistair C. K. Lam, Madeleine E. Laws, Jackson Lee, Kyle Park, Emily Wesel, Peter H. Covert, Lutz Kockel, Sangbin Park, Seung K. Kim

**Author notes:** These authors contributed equally (Anne E. Rankin, Elizabeth Fox).

## Abstract

Genome editing *in vivo* with CRISPR/Cas9 generates powerful tools to study gene regulation and function. We developed CRISPR-based methods that permitted secondary school student scientists to convert *Drosophila* GAL4 lines to LexA lines. Our novel curricula implement a new donor strain optimizing Homology-assisted CRISPR knock-in (HACK) that simplifies screening using light microscopy. Successful curricula adoption by a consortium of schools led to the generation and characterization of 16 novel LexA lines. This includes extensive comparative tissue expression analysis between the parental Gal4 and derived LexA lines. From this collaboration, we established a workflow to systematically generate LexA lines from frequently-used GAL4 lines. Modular courses developed from this effort can be tailored to specific secondary school scheduling needs, and serve as a template for science educators to innovate courses and instructional goals. Our unique collaborations highlight that resources and expertise harnessed by university-based research laboratories can transform experiential science instruction in secondary schools while addressing research needs for the community of science.

## INTRODUCTION

*Drosophila melanogaster* is a powerful organism to investigate gene function in diverse biological settings, including embryonic development and metabolism. To study genes in specific *Drosophila* organs, compartments, or cell populations, investigators have developed binary gene expression systems (Brand and Perrimon 1993; Lai and Lee 2006; Potter et al 2010; Kim et al 2021). These systems combine (1) cell-specific *cis*-regulatory elements that drive the expression of a transgene encoding an exogenous transcriptional activator (e.g., GAL4), and (2) a responder transgene whose expression is directed by the transcriptional activator. However, novel challenges in studying more complex biological contexts like inter-cellular or inter-organ communication necessitate parallel genetic manipulations of two, or more, independent cell populations. Multiple independent binary expression systems can be combined in a single fly to study genetic perturbations of multiple tissues simultaneously. This approach has led to powerful epistasis experiments between different tissues (Shim et al. 2013), simultaneous clonal lineage analysis of multiple cell populations (Lai and Lee 2006; Bosch et al. 2015), visualization of specific physical cell-cell contacts (Gordon and Scott 2009; Bosch et al. 2015, Macpherson et al. 2015), and measures of hormonal responses in target cells (Tsao et al 2022).

Simultaneous use of orthogonal binary expression systems requires generation of independent cell-specific transgenic transcriptional activators. For the LexA/LexAop binary expression system, diverse tissue-specific LexA activator lines have been systematically generated by cloning and linking putative enhancers to *LexA* (Pfeiffer et al 2010) or by inserting LexA-encoding transposons near endogenous enhancers (“enhancer trapping”; Kockel et al 2016; Kockel et al 2019; Kim et al 2023). This work enabled detailed studies of tissue-specific *LexA* expression. To expand the collection of activator lines, and to exploit the thousands of extant GAL4 lines (FlyBase) as potential targets, Lin and Potter (2016) developed Homology-assisted CRISPR knock-in (HACK) to replace GAL4 with an orthogonal transcriptional activator. Similar CRISPR/Cas9-based approaches have been successfully applied to generate LexA lines from existing GAL4 lines with well-characterized tissue expression patterns (Chang et al 2022; Karuparti et al 2023). However, no systematic approach to convert frequently used GAL4 lines to LexA has been described.

We postulated that university-based research laboratories had unrealized potential to develop and strongly influence traditional science instruction covering biology and genetics in secondary schools, and simultaneously address unmet needs - like new LexA driver lines - of the scientific community. In 2011, Stanford University investigators developed courses (hereafter, the Stan-X curricula) to introduce students to discovery-based experimental investigations based on fruit fly genetics and biology (Kockel et al 2016, 2019; Chang et al 2022; Wendler et al 2022; Kim et al 2023). Currently, there are 15 Stan-X partner schools that run semester-or year-long courses, based on ‘enhancer traping’ to generate novel LexA-expressing lines. Recently, multiple partnering schools requested diversification of Stan-X course content, including the development of ‘modular’ courses that could run for 12 weeks or less, but still provide opportunities for discovery through experimental science. In response, we developed an *in vivo* CRISPR-based curriculum with fruit flies for a consortium of schools described here. To enable brightfield microscopy-based genetic screens to score LexA flies “converted” from a GAL4 line, we developed a new yellow^+^ HACK donor strain and tested it with our secondary school consortium. Here, we report the successful generation and characterization of novel LexA lines by student scientists. This effort has established a productive paradigm of university researchers and secondary schools collaborating to generate new LexA lines based on the extensive enhancer characterization and use of its parental GAL4 line.

## MATERIALS AND METHODS

### Drosophila strains

Except for LexA.G4HACK donor lines, all other *Drosophila* lines in **Table 1**, **Figures 3**, and **S2** were obtained from the Bloomington Drosophila Stock Center (BDSC).

**Table 1.**
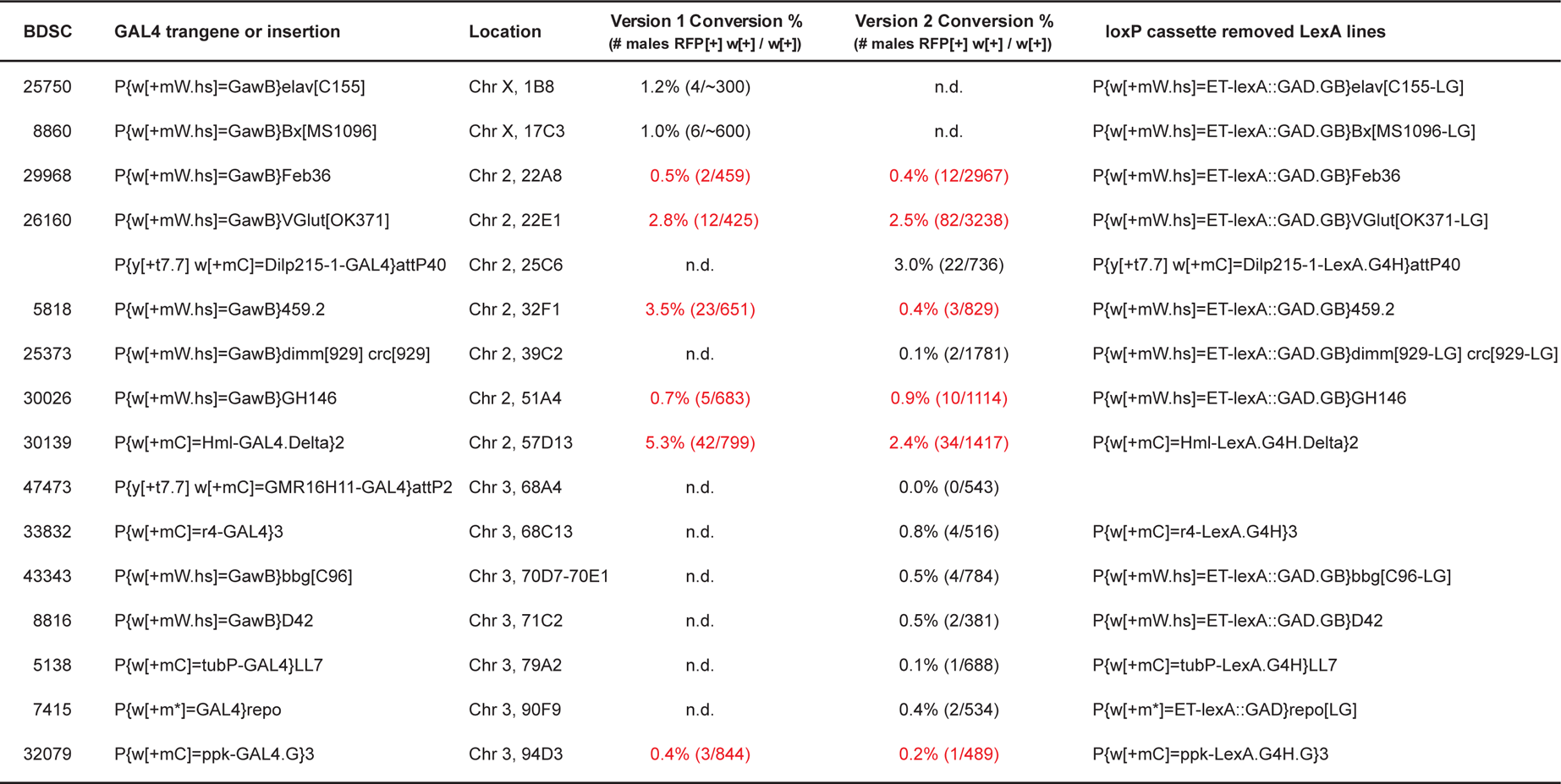
Genotypes of the original GAL4 and converted LexA lines and their conversion rate using v1 and v2 donors. Source IDs and genotypes of GAL4 lines selected for the gene conversions and their conversion rates by donor version; red color indicates lines with data for both donor versions. Following the convention of FlyBase genotype nomenclatures, converted lines from P{GawB}-based enhancer trap GAL4 insertions were named as P{ET-lexA::GAD.GB} and converted lines from cloned enhancer-driven GAL4 transgenes were named by replacing GAL4 in the original genotypes with LexA.G4H. n.d. = not determined.

| BDSC | GAL4 trangene or insertion | Location | Version 1 Conversion %<br>(# males RFP[+] w[+] / w[+]) | Version 2 Conversion %<br>(# males RFP[+] w[+] / w[+]) | loxP cassette removed LexA lines |
| --- | --- | --- | --- | --- | --- |
| 25750 | P{w[+mW.hs]=GawB}elav[C155] | Chr X, 1B8 | 1.2% (4/~300) | n.d. | P{w[+mW.hs]=ET-lexA::GAD.GB}elav[C155-LG] |
| 8860 | P{w[+mW.hs]=GawB}Bx[MS1096] | Chr X, 17C3 | 1.0% (6/~600) | n.d. | P{w[+mW.hs]=ET-lexA::GAD.GB}Bx[MS1096-LG] |
| 29968 | P{w[+mW.hs]=GawB}Feb36 | Chr 2, 22A8 | 0.5% (2/459) | 0.4% (12/2967) | P{w[+mW.hs]=ET-lexA::GAD.GB}Feb36 |
| 26160 | P{w[+mW.hs]=GawB}VGlut[OK371] | Chr 2, 22E1 | 2.8% (12/425) | 2.5% (82/3238) | P{w[+mW.hs]=ET-lexA::GAD.GB}VGlut[OK371-LG] |
|  | P{y[+t7.7] w[+mC]=Dilp215-1-GAL4}attP40 | Chr 2, 25C6 | n.d. | 3.0% (22/736) | P{y[+t7.7] w[+mC]=Dilp215-1-LexA.G4H}attP40 |
| 5818 | P{w[+mW.hs]=GawB}459.2 | Chr 2, 32F1 | 3.5% (23/651) | 0.4% (3/829) | P{w[+mW.hs]=ET-lexA::GAD.GB}459.2 |
| 25373 | P{w[+mW.hs]=GawB}dimm[929] crc[929] | Chr 2, 39C2 | n.d. | 0.1% (2/1781) | P{w[+mW.hs]=ET-lexA::GAD.GB}dimm[929-LG] crc[929-LG] |
| 30026 | P{w[+mW.hs]=GawB}GH146 | Chr 2, 51A4 | 0.7% (5/683) | 0.9% (10/1114) | P{w[+mW.hs]=ET-lexA::GAD.GB}GH146 |
| 30139 | P{w[+mC]=Hml-GAL4.Delta}2 | Chr 2, 57D13 | 5.3% (42/799) | 2.4% (34/1417) | P{w[+mC]=Hml-LexA.G4H.Delta}2 |
| 47473 | P{y[+t7.7] w[+mC]=GMR16H11-GAL4}attP2 | Chr 3, 68A4 | n.d. | 0.0% (0/543) |  |
| 33832 | P{w[+mC]=r4-GAL4}3 | Chr 3, 68C13 | n.d. | 0.8% (4/516) | P{w[+mC]=r4-LexA.G4H}3 |
| 43343 | P{w[+mW.hs]=GawB}bbg[C96] | Chr 3, 70D7-70E1 | n.d. | 0.5% (4/784) | P{w[+mW.hs]=ET-lexA::GAD.GB}bbg[C96-LG] |
| 8816 | P{w[+mW.hs]=GawB}D42 | Chr 3, 71C2 | n.d. | 0.5% (2/381) | P{w[+mW.hs]=ET-lexA::GAD.GB}D42 |
| 5138 | P{w[+mC]=tubP-GAL4}LL7 | Chr 3, 79A2 | n.d. | 0.1% (1/688) | P{w[+mC]=tubP-LexA.G4H}LL7 |
| 7415 | P{w[+m*]=GAL4}repo | Chr 3, 90F9 | n.d. | 0.4% (2/534) | P{w[+m*]=ET-lexA::GAD}repo[LG] |
| 32079 | P{w[+mC]=ppk-GAL4.G}3 | Chr 3, 94D3 | 0.4% (3/844) | 0.2% (1/489) | P{w[+mC]=ppk-LexA.G4H.G}3 |

### Generation of version 1 and version 2 LexA.G4HACK donor strains

The construction of pHACK-GAL4>nlsLexA::GADfl (v1) and its insertion into PBac{y+-attP-9A}42A13 on the CyO balancer chromosome were described previously (Chang et al 2022). The CyO balancer chromosome with the v1 donor transgene was combined with the PBac{y[+mDint2] GFP[E.3xP3]=vas-Cas9}VK00027 transgene on the third chromosome (BDSC 51324) to make a fully functional v1 donor strain as previously reported (Chang et al 2022) 4965 bp PCR fragment was amplified from pCaryP (Groth et al 2004) using the primers y[+]_F2 (5’-ATTAGTCTCTAATTGAATGACGTCGCATACTTACATTTTTTCCGCTTTTTCCG-3’) and y[+]_R (5’-GCTATACGAAGTTATGACGTCGTCGACTATTAAATGATTATCGCCCGATTACC-3’) and inserted to AatII site on pHACK-GAL4>nlsLexA::GADfl (v1) using the HiFi DNA Assembly Cloning Kit (New England BioLabs, E5520S). The resulting construct carrying both 3xP3-RFP and *yellow* transgene markers, pHACKy-GAL4>nlsLexA::GADfl (v2), was inserted into the PBac{y+=attP-9A}42A13 site on the CyO chromosome (the same site as the v1 donor construct). The CyO balancer chromosome with the v2 donor transgene was combined with the M{GFP[E.3xP3]=vas-Cas9.RFP-}ZH-2A transgene on X chromosome (BDSC 55821) to make a fully functional v2 donor strain.

### Intercross strategy for CRISPR/Cas9-based conversion of GAL4 to LexA.G4HACK

For the F0 intercross, each vial contained four males of the GAL4 line and four virgin females of the LexA donor line (either v1 or v2). The F0 intercross was transferred to new vials every three days for two weeks. When F1 progeny emerged, each male progeny carrying w[+] and CyO was mated to two virgin females of y[1] w[1118] (BDSC 6598). At least 20 mating pairs were set up to identify independent conversion events from different males. These F1 mating pairs were transferred to new vials once after 5 days of mating to extend the number of F2 male progeny to screen for, but we found this may not be necessary if 40 or more mating pairs were initially set up. For the v1 HACK donor line, F2 male progeny with w[+] and non-CyO markers were selected and screened for RFP expression in ocelli under a fluorescence stereo microscope. For the v2 HACK donor line, we screened for males carrying w[+], y[+], and non-CyO markers under a light stereo microscope, then confirmed their RFP expression in ocelli under a fluorescence stereo microscope. All F2 male progeny with w[+] and non-CyO markers were counted to calculate the overall conversion rates in **Table 1**. To assess the HACK-mediated gene conversion efficiency in independent male germlines, we measured frequencies of gene conversion events from each mating pair and plotted them in **Figure S1**. GAL4 stocks usually carry a wild-type Y chromosome, but we have noted that some GAL4 stocks harbor undocumented Dp(1;Y)y^+^ chromosomes, and could interfere with body color-based screening in the F2 generation. Two independently converted males per each GAL4 line were saved for further analysis.

### Removal of loxP cassette from HACK-converted LexA.G4 lines

A single converted F2 male was mated to two virgin females carrying P{Crey} on the X chromosome (BDSC 766). A single F3 male carrying the w[+] marker was mated to two virgin females of y[1] w[1118] (BDSC 6598). A single founder F4 male with w[+], but without the y[+] cuticle color marker or RFP expression in the ocelli was mated to a balancer line (e.g. BDSC 59967) to isolate the chromosome carrying LexA.G4H with only w[+] marker. Even without a heat shock, all F4 males that we have seen were without RFP and y[+] markers, indicating high expression of Cre in F3 male germlines harboring the P{Crey} transgene.

### PCR genotyping and sequencing of converted LexA.G4 lines

Genomic DNAs from the original GAL4, HACK donor, and converted LexA male flies were extracted as previously reported (Chang et al 2022). 1 μl of the extracted genomic DNA was added to 19 μl of PCR master mix containing 7 μl of water, 10 μl of Q5 Hot Start High-Fidelity 2x Master Mix (NEB M0494S), 1 μl of 10 μM Primer 1 (5’-ATGAAGCTACTGTCTTCTATCGAACAAGC-3’) for a GAL4 sequence, and 1 μl of 10 μM Primer 2 (5’-GGCATACCCGTTTGGGATATATGATCC-3’) for a HACK donor sequence. After a 30-second denaturing period at 98°C, 35 cycles of PCR amplification were performed as a 10-second denaturing period at 98°C, a 30-second annealing period at 60°C, and a 1-minute extension period at 72°C. The PCR reactions from GAL4, donor, and converted flies were resolved in TAE-agarose gel electrophoresis. 1367 bp-long PCR product was amplified only from converted flies, isolated using Zymoclean Gel DNA Recovery Kit (Zymo Research D4008), and sequenced from both ends using Primer 1 and Primer 2.

### Imaging LexAop-GFP reporter gene expression

P{10XUAS-IVS-mCD8::GFP}attP2 (BDSC 32185) and P{13XLexAop2-mCD8::GFP}attP2 (BDSC 32203) were used to compare the expression patterns of the original GAL4 and converted LexA.G4HACK line pairs. Four virgin females carrying GFP reporters were mated to a single male of either GAL4, LexA.G4H (RFP^+^), or LexA.G4H (RFP^−^) lines. The mating pairs were transferred to new vials every two days until imaging of expression patterns had been completed. For imaging larval tissues, inverted third instar larvae at the wandering stage were fixed at 4% paraformaldehyde in PBS for >16 hours at 4°C and washed three times in PBS containing 0.1% Triton X-100. Larval brains and imaginal discs were dissected from the washed carcass, transferred onto a glass slide, immersed in 6 μl of the mounting media with DAPI (Vectashield H-1200) for 1 minute, and mounted under an 18 x 18 cover glass. Images of GFP, RFP, and DAPI channels were captured on a compound fluorescence microscope and edited using ImageJ software (NIH). For live imaging of early pupal hemocytes, third instar larvae at the wondering stage were starved on a 2% agar plate for 4 hours, and circulating hemocytes in pupating larvae were imaged under a fluorescence stereo microscope for 30 seconds (Movie S1 and https://youtu.be/Bk EaKTiVE).

## RESULTS

### A simplified genetic strategy for identifying successful gene conversion *in vivo*

A red fluorescent eye marker, 3xP3-RFP, was used in the original HACK study to detect successful editing of GAL4 (Lin and Potter 2016, Chang et al 2023; hereafter the version 1 donor, or “v1”). However, genetic screening requiring fluorescence microscopy could prevent adoption of HACK by schools with limited budgets. To permit screening for successful HACK gene conversion with light microscopy, we produced a new transgenic donor strain harboring a 5 kb transgene carrying the *yellow* gene enhancer and intronless coding sequence, inserted next to 3xP3-RFP transgene (**Figure 1A**: see Methods). Briefly, we generated a plasmid construct called pHACKy-GAL4>nlsLexA::GADfl (version 2 donor or “v2” hereafter) and inserted this in the attP42A13 genomic site on the CyO balancer, the same position as pHACK in v1 donors (**Figure 1A**: Methods). Unexpectedly, adults harboring the v2 donor had enhanced RFP expression in eyes and ocelli compared to the v1 donor at the same molecular location (**Figure 1B**), indicating that the 5 kb *yellow* transgene may have improved the expression of the neighboring 3xP3-RFP transgene in this genomic location.

**Figure 1.**
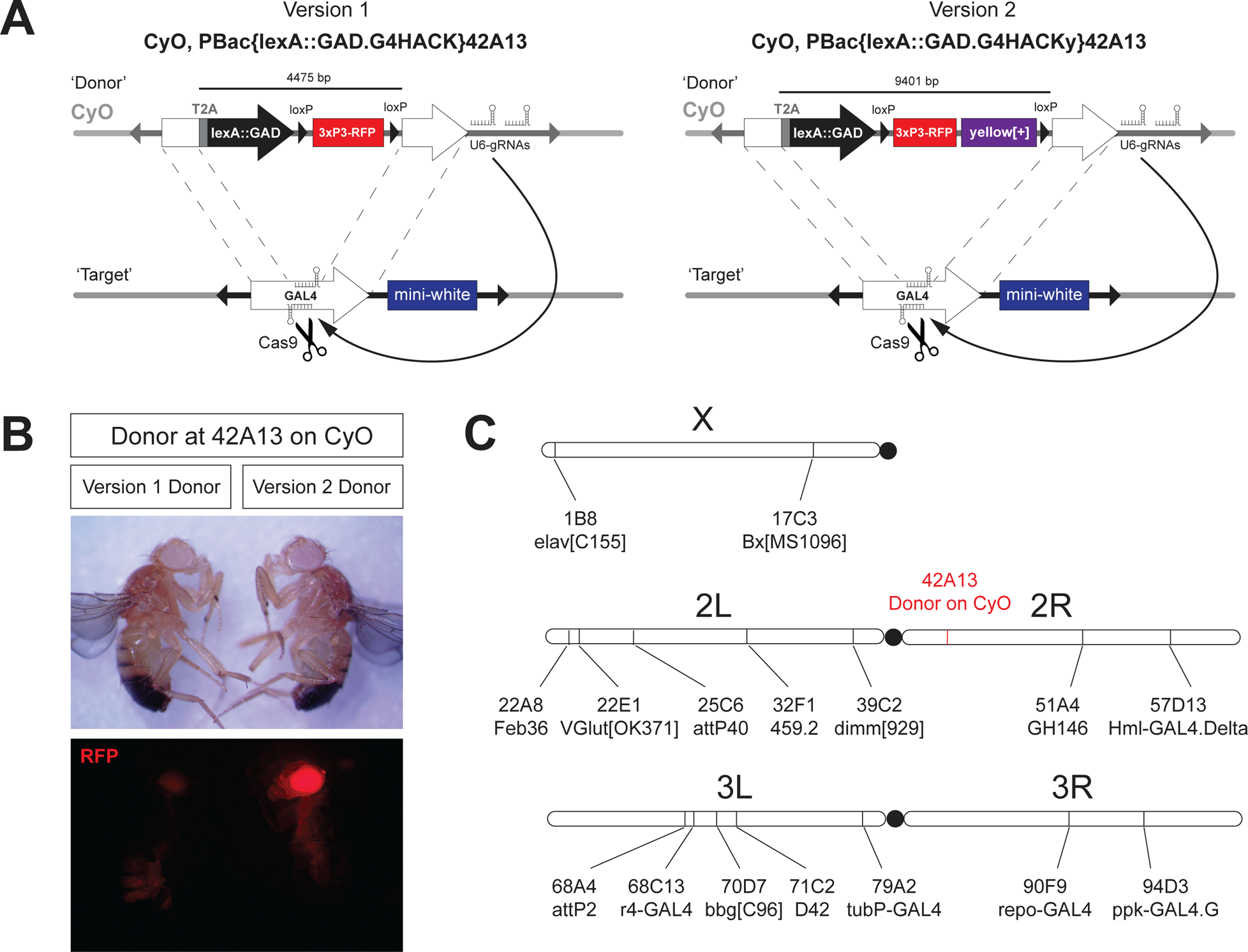
Designs of LexA.G4HACK donors for CRISPR/Cas9-mediated GAL4 gene conversion and chromosomal locations of GAL4 targets. (A) Genetic designs of two LexA.G4HACK donors for HACK-mediated gene conversion. A DNA double-strand break generated by vas-Cas9 and gRNAs targeting the GAL4 sequence in germline chromosomes can be repaired by homology-assisted CRISPR/Cas9 knock-in of a donor transgene located in a balancer chromosome. The version 2 donor carries a loxP-flanked dual transgene cassette. Both versions of the donor are inserted in the same attP site on the CyO balancer to enable an unbiased comparison of the donor efficiency differences potentially generated by different repair template sizes. (B) The version 2 donor transgene at the genomic location of 42A13 on the CyO balancer showed an improved 3xP3-RFP expression compared to the version 1 donor at the same location. *yellow*^+^ phenotypes in both flies shown are from PBac{y+-attP-9A}42A13 on the CyO balancer. (C) Chromosomal locations of selected GAL4 targets for HACK-mediated gene conversion and the donor location

To determine whether the additional 5 kb payload in the v2 donor and the use of a different *Cas9* transgene would affect the overall HACK efficiency, we measured GAL4>LexA.G4H conversion in six GAL4 lines (red font in **Table 1**) using v1 and v2 HACK donors. For the v1 donor experiment, the *PBac{vas-Cas9}VK00027* transgene located on the third chromosome (BDSC 51324: Port et al 2015) was used (Chang et al 2022). With the v2 donor, we switched to the X-linked *M{vas-Cas9.RFP-}ZH-2A* transgene in a *yellow* background (BDSC 55821, Port et al 2015) to facilitate screening of *yellow* transgene integration events (**Methods**). Overall HACK efficiencies of v2 were slightly lower (1.4%, n = 10,054) than those of the v1 donor (2.3%, n = 3,861). However, the relative HACK efficiencies among the different target locations appeared similar between v1 and v2 except for the 32F1 location, indicating that the v2 HACK donor is comparable to v1 in GAL4 target-gene conversion efficiency.

To assess the frequency of gene conversion (GAL4 to LexA) in the germ cell lineage of individual male flies, we measured the frequencies of conversion events stemming from individual male matings. This was contrasted with the measurement of the overall conversion rate (**Table 1**), which reflects data pooled from a standard-sized F1 intercross (n = 40); this quantification scheme differs slightly from a prior study (Lin and Potter, 2016), which combined data from 4 males to determine conversion rates. Conversion frequency from an individual F1 male was scored (red number on each bar in **Figure S1**). In the lines with higher overall conversion rates (OK371-GAL4 and Hml-GAL4 in **Figure S1**), we observed that only a few male germ lines (12/94 and 3/59) produced a large number (3 or more) of conversion events, while conversions were more frequent from independent males (40/94 and 17/59). Conversely, lines with lower overall conversion rates produced conversions less often from independent males (2/23 for 459.2-GAL4 and 2/37 for dimm-GAL4), but did not necessarily produce a smaller batch of conversion events (all 22 events found in 1/16 mating for Ilp215-1-GAL4 at attP40). Our finding that conversion events occur “infrequently in many crosses” rather than “frequently in a few crosses” suggests that parallel screening of a relatively large number (e.g., n = 40) of germlines would be the more efficient approach rather than a serial screening of a smaller number (e.g., n = 20) of germlines (**Methods**).

HACK-mediated gene conversions on second chromosome-linked GAL4 lines (*cis*-chromosomal HACK) were all successful (n = 7/7), with efficiencies averaging between 0.1 and 5.3% (**Table 1**). For third chromosome-linked GAL4 lines (*trans*-chromosomal HACK) 6/7 conversions were successful, but the average conversion efficiency was lower (0 to 0.8%: **Table 1**). Prior studies of *cis*-chromosomal HACK found that HACK donors more proximal to *cis*-targets converted at higher efficiency than distal donors (Lin and Potter, 2016). However, with v1 or v2 HACK donors on the CyO balancer second chromosome, we did not observe this proximity effect on two homologous chromosomes. For example, using distally-located (42A13) donors on the CyO balancer, two GAL4 targets closely located at 22A8 and 22E1 show respective HACK efficiencies of 0.4-0.5% versus 2.5-2.8%. Thus, gene conversion efficiencies for GAL4 insertions did not reflect the mere chromosomal distance between the donor and target in *cis*-chromosomal HACK, even when compensating for inversions and translocations on the CyO balancer chromosome. This further supports the strategy of using a single HACK donor embedded in a balancer chromosome, even for *trans*-chromosomal conversions, rather than providing specific HACK donors closely located to a particular GAL4 position for each *cis*-chromosomal HACK. In sum, the v2 HACK donor on the CyO balancer showed comparable performance to v1 and can be used for both *cis-* and *trans-*chromosomal HACKing of GAL4 lines to LexA.G4H.

### Visible phenotypes permit the detection of successful HACKing

Based on our observation of brighter RFP expression in v2 donor flies compared to v1 donors (**Figure 1B**) we postulated that this difference might persist after CRISPR-based GAL4>LexA.G4H conversion. We compared RFP expression after conversion at four different genomic locations (22A8, 22E1, 68C13, and 94D3). In each, the integrated v2 donor showed bright RFP expression in ocelli (arrows, **Figure 2A**). To assess RFP expression after CRISPR/Cas9-mediated targeting with v2 donors at diverse genomic target locations, we compared heads of ten converted GAL4>LexA.G4H flies (**Figure 2B**). After the successful conversion of all ten lines, we observed that RFP expression in compound eyes was variable at different loci, as previously reported (Horn et al 2000), but RFP expression in ocelli cells was observed in all integration sites. Thus ocelli-based screening provides a reliable method for identifying v2 donor-generated conversion events with a fluorescence stereomicroscope.

**Figure 2.**
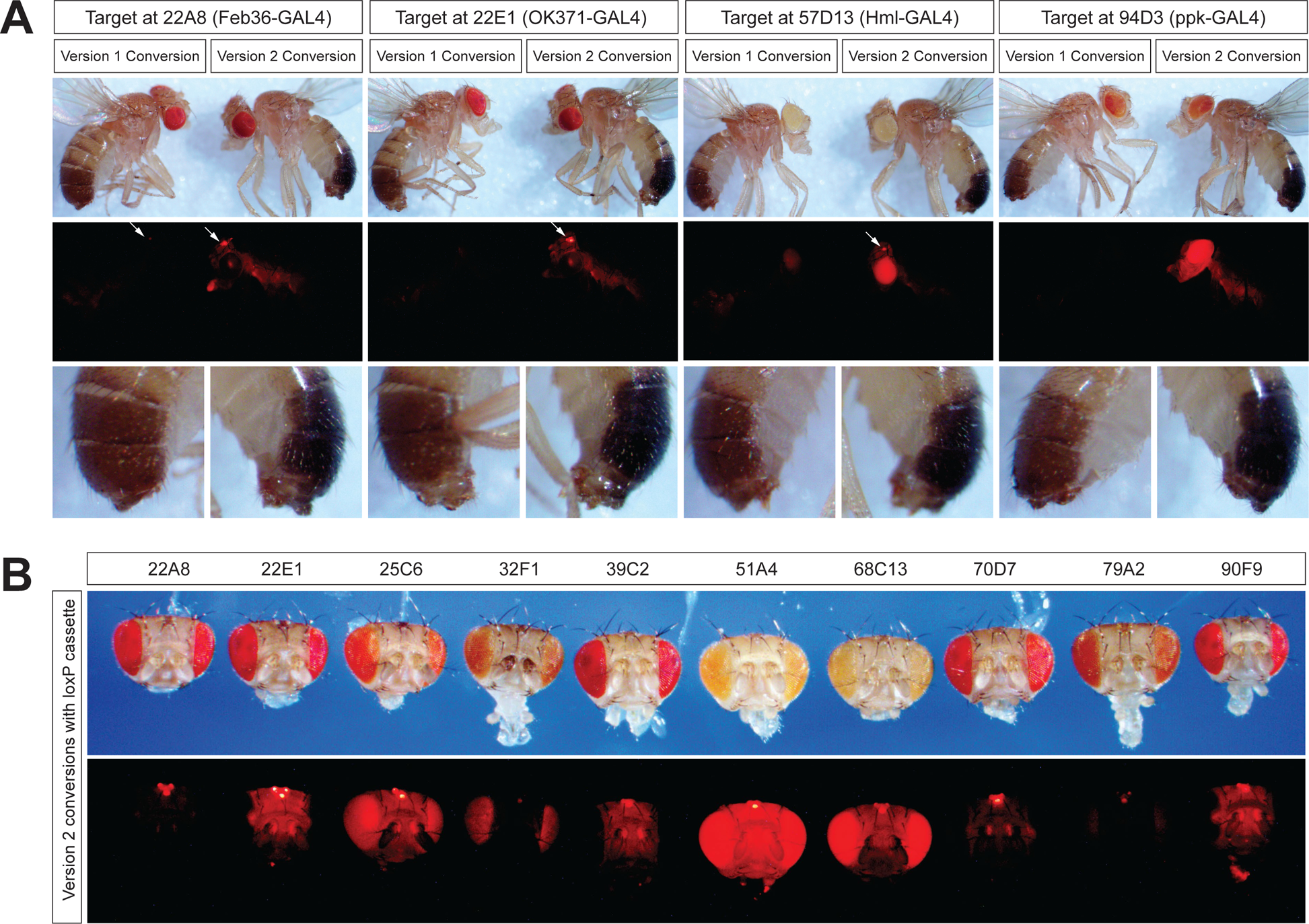
Improved RFP expression of integrated version 2 donor at various genomic locations. (A) Phenotypic comparison of F2 males with successful donor integrations at different targets. RFP expression in ocelli (white arrows) was more consistently observed in version 2 integration sites than in corresponding version 1 integration sites. The version 2 integration events can also be identified by yellow transgene expression in tail segments in the *y*^1^ *w*^1118^ mutant genetic background (the bottom row). (B) RFP expression of integrated version 2 donor at different genomic locations. Adult heads of converted males were arranged based on target locations. RFP expression in ocelli was consistently high in all locations, but the expression in compound eyes was highly variable in different locations. Note that the expression of mini-white and 3xP3-RFP was inversely correlated in compound eyes (see text).

In addition to RFP expression, conversion with the v2 donor also led to progeny with visibly darker pigmented abdominal segments, consistent with expression of the *yellow* transgene (*y*^+^) in a *yellow* mutant (*y*^1^) genetic background (**Figure 2A**). Thus, the yellow transgene embedded in the v2 donor sequence simplified screening for HACKy-mediated gene conversion events with bright-field microscopy (**Figure 3**). In summary, two markers in the v2 donor - *yellow* and RFP - facilitated screening of HACKy-mediated gene conversion events using light or fluorescence microscopy.

**Figure 3.**
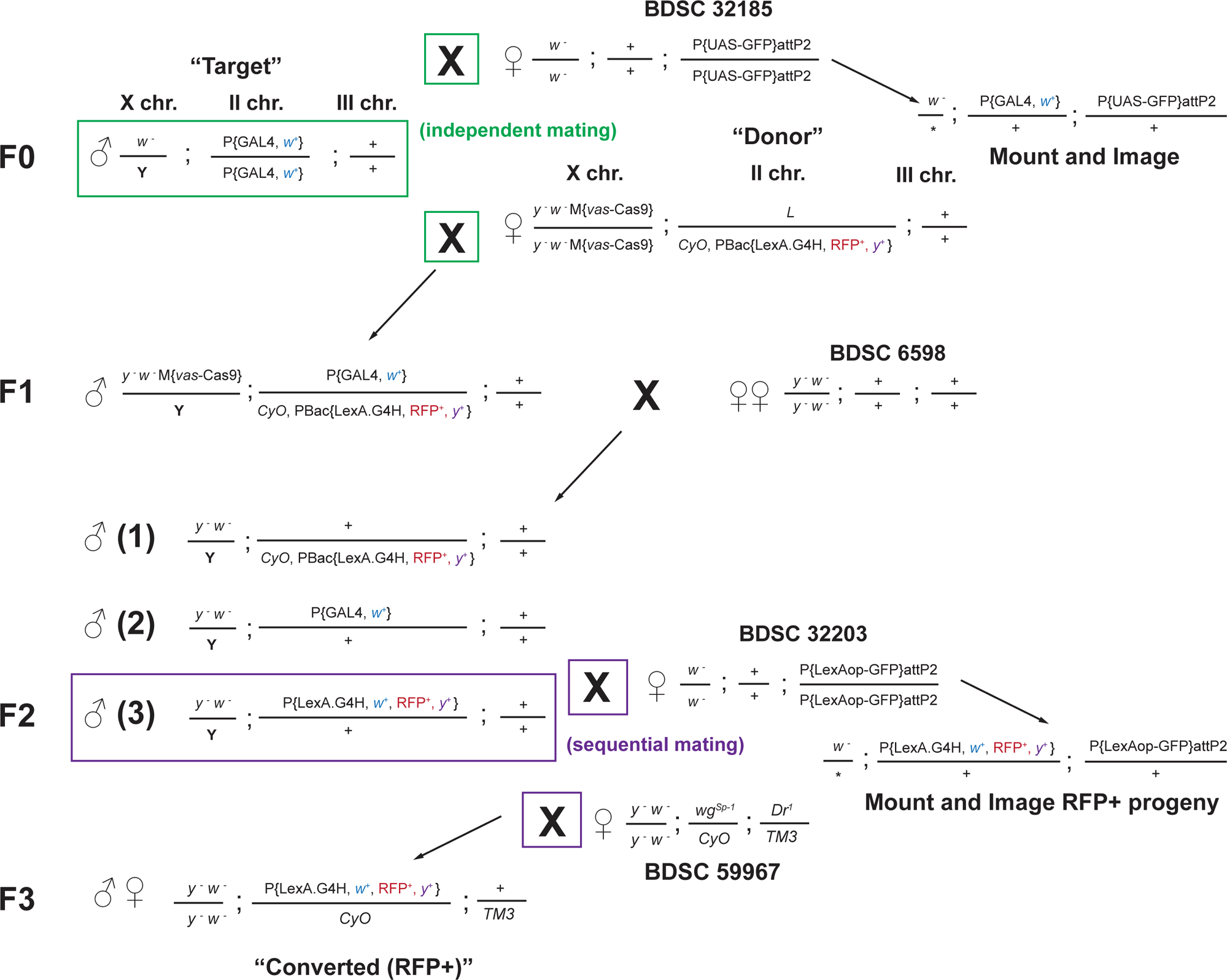
Mating scheme for converting second chromosome-linked GAL4 lines to LexA.G4H and imaging reporter expression. The parental mating (F0) was set up with a male carrying the GAL4 transgene (“Target”) and virgin females carrying vasa-Cas9 on the X chromosome and a HACK donor on the CyO balancer (“Donor”). In parallel, a male carrying the same “Target” GAL4 transgene was also mated with virgin females carrying the UAS-GFP transgene (BDSC 32185) for documentation of the GFP expression pattern of the “Target” GAL4. These mating pairs were transferred to new vials every two days 6 times. Larval, pupal, and adult progeny from UAS-GFP mating were imaged for GFP expression patterns based on prior characterizations of the “Target” GAL4 line. Up to 80 individual mating pairs were set up for an F1 male progeny carrying all three transgenes and two virgin females of *y*^1^ *w*^1118^ (BDSC 6598). In the F2 generation, non-curly male flies carrying mini-white transgene were scored for RFP expression in ocelli and/or yellow transgene expression in tail segments. If identified, a single F2 male carrying the mini-white and RFP transgenes was first mated with virgin females carrying LexAop-GFP transgene (BDSC 32203) for 3 days. The same male was mated again with different virgin females carrying balancer chromosomes (e.g., BDSC 59967) to isolate the chromosome with the modified transgene (“Converted (RFP^+^)”).

### Tissue expression patterns of originating GAL4 and converted LexA lines

To test if LexA expression in converted lines was identical to that in the originating GAL4 line, we performed intercrosses to assess and compare LexA-dependent and GAL4-dependent reporter gene expression. A single male from each original GAL4 line was mated to virgin females carrying 10xUAS-mCD8::GFP, and a single converted LexA.G4H male from each screen was mated to virgin females carrying 13xLexAop2-mCD8::GFP (**Figure 3**). To minimize the positional effects of reporter transgene expression, we used GFP reporter transgenes located at the same genomic location on the third chromosome, attP2 (Pfeiffer et al 2010).

The expression patterns of GFP in the converted LexA lines matched that of the original GAL4 lines (**Figure 4A and B**), an assessment that was less ambiguous after Cre-mediated excision of the donor *loxP*-flanked 3xP3-RFP and *yellow* transgene cassettes (see RFP^+^ cells in **Figure 4A and B**; **Figure S2**). While this *loxP*-flanked transgene cassette did not appear to alter the expression of LexA lines, we removed this cassette in all converted LexA lines.

**Figure 4.**
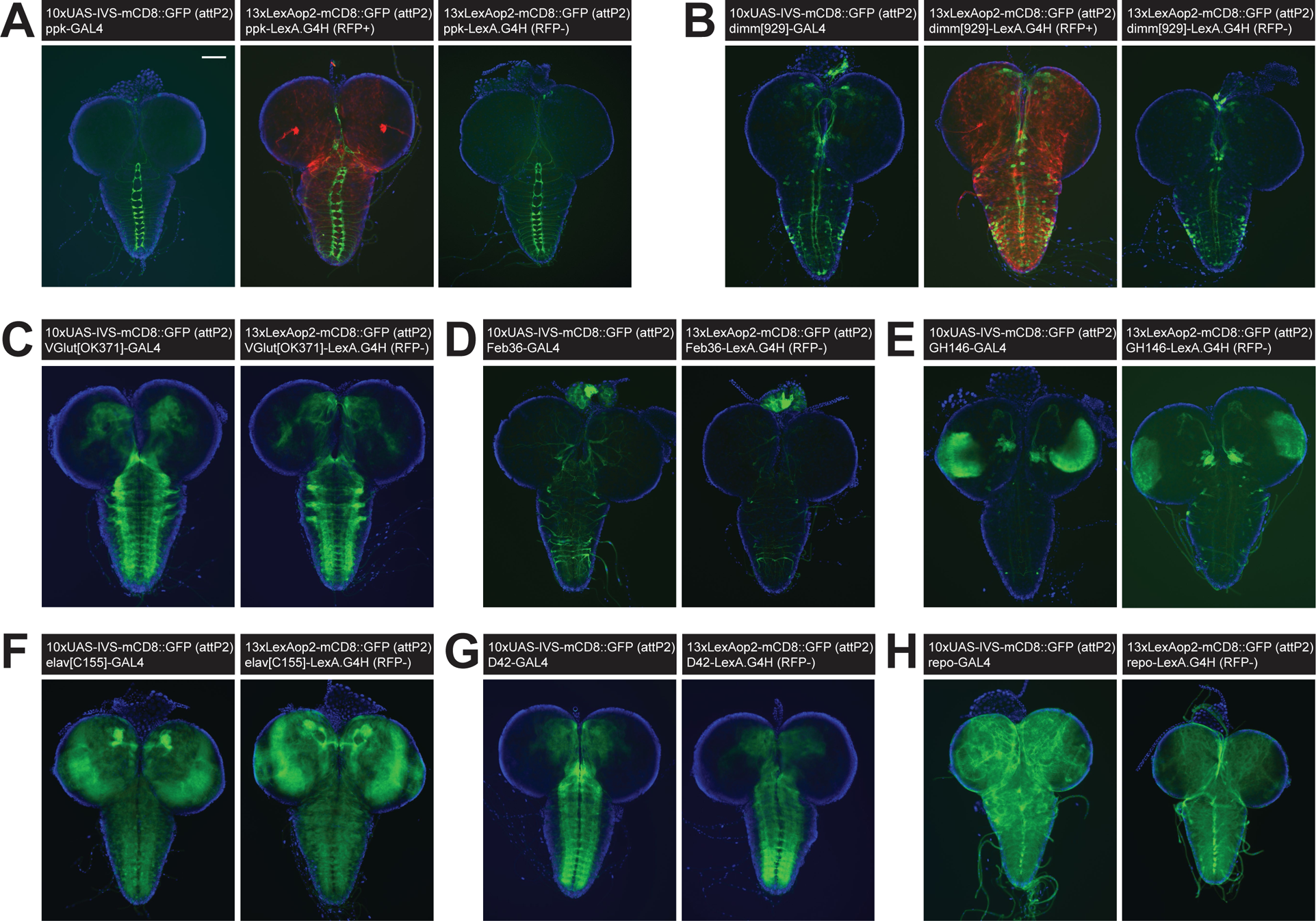
Comparison of larval brain reporter expression for original GAL4 and converted LexA.G4H lines. (A) GFP reporter expression in ventral nerve cords of larval brains driven by ppk-GAL4 (left), ppk-LexA.G4H with RFP transgene (middle), and ppk-LexA.G4H with RFP cassette removed (right). The scale bar is 100 μm. (B) GFP reporter expression in neuroendocrine cells of larval brains driven by dimm-GAL4 (left), dimm-LexA.G4H with RFP transgene (middle), and dimm-LexA.G4H with RFP cassette removed (right). (C-H) GFP reporter expression in larval brains driven by GAL4 (left) and LexA.G4H with RFP cassette removed (right) showing expression in vGlut neurons by OK371 enhancer (C), corpora cardiaca cells by Feb36 enhancer (D), brain hemispheres by GH146 enhancer (E), pan-neuronal cells by C155 enhancer (F), ventral nerve cords by D42 enhancer (G), and pan-glial cells by a cloned repo enhancer (H).

In the third instar larval brains of converted LexA.G4H lines, GFP reporter expression patterns appeared indistinguishable from reporter expression in the original GAL4 lines (**Figure 4**). However, the intensity of GFP signal of some converted LexA lines (**Figure 4E and H**) appeared slightly reduced, compared to reporter GFP signal in the original GAL4 lines. In the third instar wing discs, the converted LexA lines that drive reporter expression in the dorsal compartment of the wing disc (**Figure 5A**), the entire wing disc (**Figure 5B**), or the dorso-ventral boundary of the wing disc (**Figure 5C**) all showed identical patterns to the original GAL4 lines. In whole animal live imaging, mCD8::GFP signals on circulating hemocytes that migrate from anterior to posterior in early pupa (Movie S1; **Methods**) also appeared identical between GAL4 and LexA lines (**Figure 5D**). Compared to the original GAL4 lines, converted lines expressing LexA in the adult abdomen and head fat body also showed similar reporter GFP expression patterns (**Figure 5E**). Taken together, our analysis confirmed that the transactivation functions of converted LexA.G4H lines are indistinguishable from the original GAL4 fly lines.

**Figure 5.**
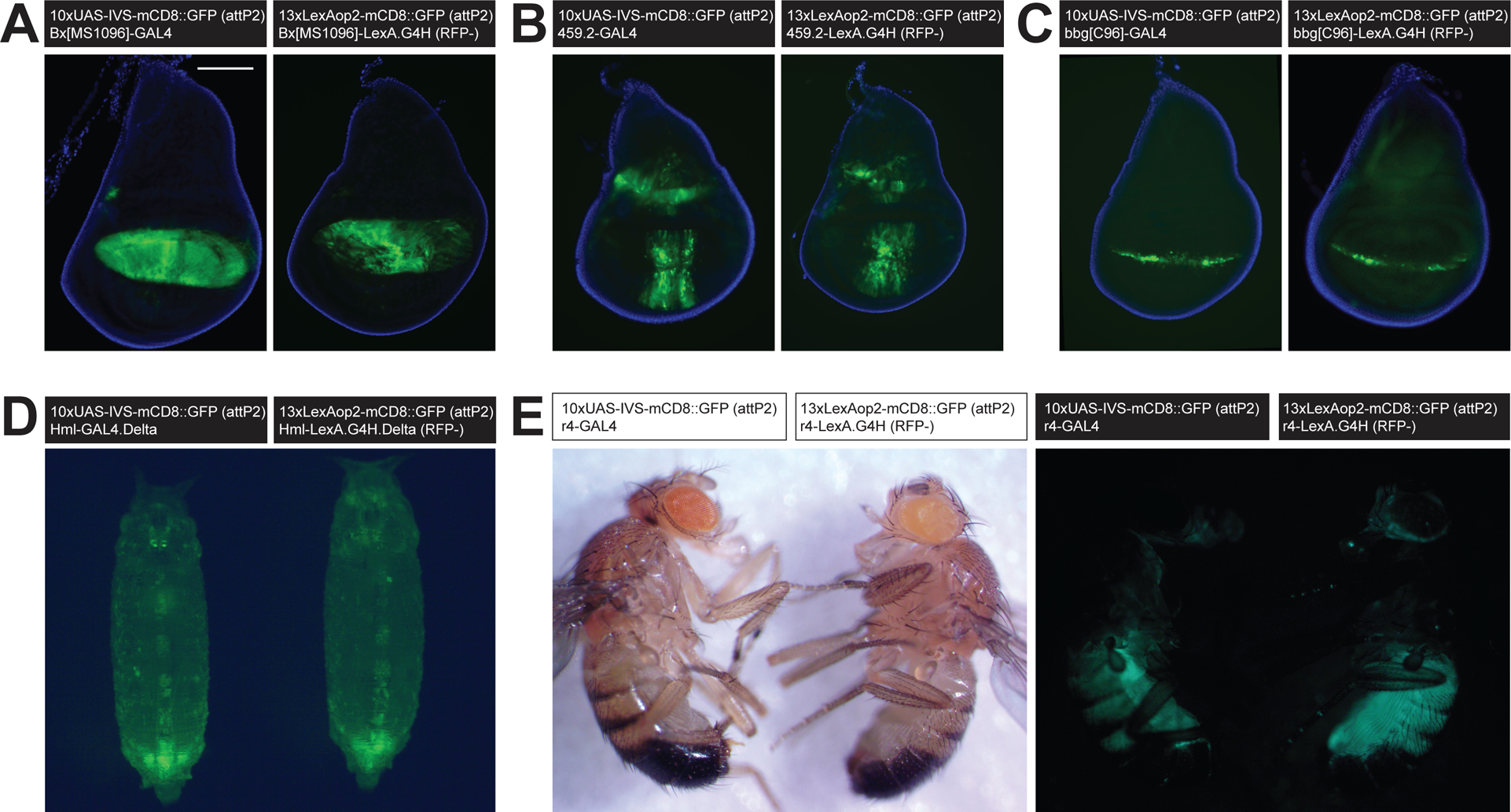
Comparison of reporter expression for original GAL4 and converted LexA.G4H lines in larval wing discs, pupal hemocytes, and adult fat bodies. (A-C) GFP reporter expression in larval wing discs driven by GAL4 (left) and LexA.G4H with RFP cassette removed (right) marking cells in the dorsal pouch by MS1096 enhancer (A), broad anterior-posterior boundaries by 459.2 enhancer (B), and dorsal-ventral boundaries by C96 enhancer (C). The scale bar in (A) is 100 μm. (D) GFP reporter expression in early pupae driven by GAL4 (left) and LexA.G4H with RFP cassette removed (right) showing expression in circulating hemocytes by a cloned Hml enhancer. The image is a still frame from 30-second-long live imaging (Movie S1), (E) GFP reporter expression in the fat body of adult males driven by r4-GAL4 (left side of each image) and r4-LexA.G4H with RFP cassette removed (right side of each image).

### Innovating secondary school curricula for systematic generation of LexA enhancer lines

To develop science classes that used CRISPR/Cas9 mediated gene conversion to generate novel LexA-expressing fly lines, we partnered with secondary schools that had previously collaborated with us to develop relevant fruit fly-based science instruction curricula (**Methods**: https://www.stan-x.org). As conversion targets, we selected six GAL4 lines whose expression patterns were previously well-characterized. We sequentially developed two courses for teaching fly genetics covering CRISPR/Cas9-mediated gene conversion, larval tissue dissection, and fluorescence imaging techniques (**Figure 6**). The first course (2021) used the v1 donor, while the following class (2022) used donor v2.

**Figure 6.**
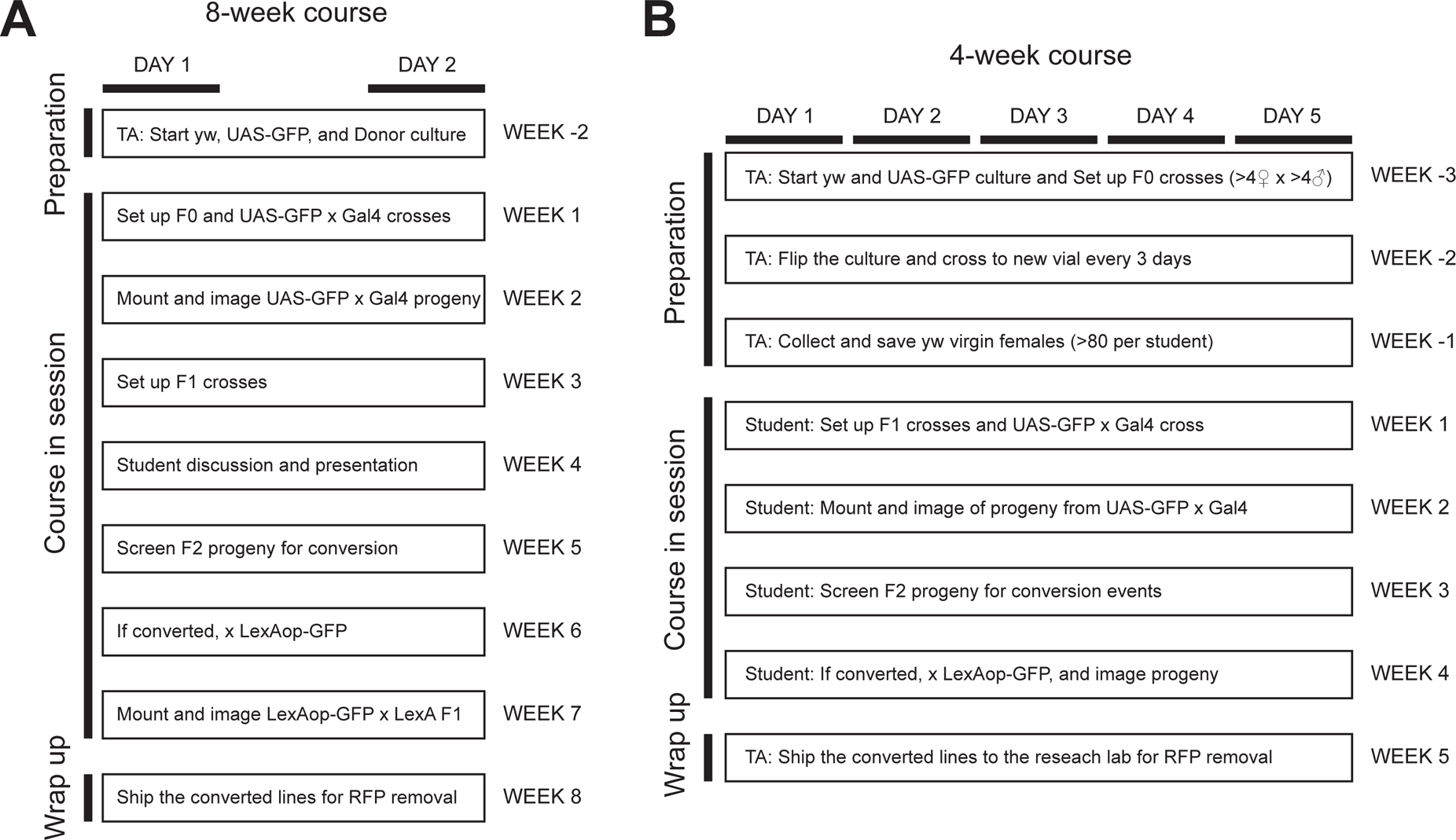
Genetics laboratory class schedules deployed for 8 weeks and 4 weeks in secondary schools. (A) A 90-minute-long class was held twice a week for 8 weeks. In week 1, students were introduced to *Drosophila* genetics including understanding genotypes, identifying associated markers, and setting up mating with follow-up maintenance. In week 2, students learned the anatomy of the third instar larva, micro-dissection, and imaging of slide-mounted tissues. In weeks 3 and 4, students generated a series of F1 intercrosses, while participating in discussions of prior characterizations of assigned GAL4 lines. In weeks 5 and 6, students started screening for conversion events in F2 progeny. In week 7, if a converted male was found, they set up a mating with LexAop-GFP reporter and documented GFP expression in the resulting progeny with RFP. In week 8, instructors and teaching assistants shipped the “converted (RFP^+^)” lines to research laboratories. (B) A 6-hour-long daily class was held five days a week for 4 weeks. To start week 1 with F1 mating, instructors and teaching assistants initiated F0 mating 3 weeks before the class started while collecting and maintaining virgin females of *y*^1^ *w*^1118^. Since the daily class schedule permitted students to master micro-dissection and imaging techniques more thoroughly, all students successfully documented GFP expression by assigned GAL4 lines by the end of week 2. Students had hour-long daily remote meetings with a research scientist to troubleshoot and discuss primary research articles.

In the first course using the v1 donor, six students focused on experimental design, execution, and interpretation, and successfully converted assigned GAL4 lines (red font in **Table 1**) over a 10.5 week schedule (Phillips Exeter Academy, NH; **Figure 6A**). Students performed intercrosses and screened for a “HACKed GAL4”, then stabilized the chromosome carrying each converted LexA driver over a balancer chromosome. Subsequently, the class imaged and compared LexA.G4H-dependent GFP reporter expression with the original GAL4-dependent expression in larval tissues. The class performed genomic PCR and sequencing to genotype converted lines (confirming substitution of LexA.G4H for GAL4: **Methods**), but prioritized functional verification of the converted LexA line by GFP reporter expression. Other class time was devoted to understanding prior usage of the original GAL4 lines. Suggestions from students and instructors for improving the course workflow included: (1) enhancing RFP expression in future studies to ease screening and identification of converted LexA lines, and (2) considering additional visible phenotypes to identify converted flies, since access to fluorescence stereomicroscope during this course was a significant ‘bottleneck’.

To address these suggestions, we developed the v2 donor and tested its use in a second course (The Lawrenceville School, NJ; **Figure 6B**). To perform two generations of mating within a 4.5 week summer schedule, an instructor and a teaching assistant intercrossed F0 mating pairs two weeks before the course commencement: this allowed students on the first day of class to identify the correct F1 male progeny, and to set up intercrosses with virgin female flies (**Methods**). Despite the demanding schedule, 5/6 students identified at least one LexA.G4H convertant. Removal of the loxP-RFP cassette was performed separately by a university research partner (**Figure S2**). This subsequent work (1) established balanced, ‘genetically stable’ LexA lines in a uniform genetic background (*y*^1^ *w*^1118^), (2) verified LexA.G4H-dependent tissue expression of a GFP reporter, and (3) distributed new lines to a *Drosophila* stock center, which makes stocks available for an international community of scientists. In summary, these interscholastic curricula and collaborations established CRISPR/Cas9-based strategies to generate well-characterized, novel LexA fruit fly lines ready for use by the science community. Additionally, both classes provided ‘proof of concept’ for the feasibility of applying an *in vivo* CRISPR/Cas9 genome editing curriculum in a secondary school setting.

## DISCUSSION

To expand the collection of LexA drivers, we and others have generated novel LexA lines using enhancer trap screens (Kockel et al 2016, Kockel et al 2019, Kim et al 2023) or by cloning enhancers to direct LexA expression (Pfeiffer et al 2010; Wendler et al 2022). While these approaches are sound, the novel lines generated by random transposon insertion or putative genomic enhancer fragments require extensive characterization, including insertion site mapping or expression specificity. As an alternative, complementary approach, CRISPR/Cas9 ‘HACK’ strategies to generate LexA lines that recapitulate tissue expression patterns of existing GAL4 lines were recently developed. We have modified these approaches (Lin and Potter 2016; Chang et al 2022) to generate new LexA lines, substantially simplifying the screening of HACK events using visible body color phenotypes (HACKy). GAL4>LexA.G4H gene conversion can be subsequently confirmed by detecting eye/ocelli expression of a second RFP marker. In multiple cases, we observed identical tissue expression patterns of reporter genes induced by the original GAL4 and the cognate converted LexA.G4H line, demonstrating the high fidelity of HACKy-mediated conversion. To address demands for experiential science instruction, we worked with secondary school partners to develop curricula that systematically generated new LexA lines with well-characterized gene expression patterns. GAL4 lines were prioritized based on the characterization of the desired expression, and frequency of cited usage (http://flybase.org/GAL4/freq_used_drivers/). Our work with student scientists demonstrates how university-based research could be leveraged to achieve educational outreach that also generates useful tools for the community of science.

Using the second chromosome-based v2 donor, gene conversion efficiencies of second chromosome-linked GAL4 lines were higher on average than those observed with third chromosome-linked GAL4 lines. This indicates that *cis*-chromosomal HACKy remains more efficient than *trans*-chromosomal HACKy. Thus, additional lines to achieve *cis*-chromosomal HACKy of third chromosome-linked GAL4 lines could be useful.

Prior studies showed that most non-converted F2 males contain small deletions at target GAL4 sequences, indicating the prevalence of non-homologous end joining repair during HACK (Lin and Potter 2016). Thus, we speculate that after CRISPR/Cas9 DNA targeting, biasing homology-directed repair over non-homologous end joining (NHEJ) at double strand breaks could improve conversion efficiency. One possibility to achieve this would be to construct donor strains with impaired NHEJ (Beumer et al 2013).

Recent exciting advances in biology, like CRISPR gene editing, provide opportunities for secondary school instructors to refresh and invigorate curricula targeting nascent student scientists. To leverage this progress, we developed an experimental curriculum that: (1) incorporated several vibrant areas of bioscience, including genetics, molecular biology, bio-informatics, developmental biology, and evolutionary biology, (2) centered around a powerful modern gene editing technology (CRISPR/Cas9 and homology-directed repair) widely-known to the general population that captured the interest of students and their instructors, (3) was based in fruit flies, a cost-effective, safe experimental system with rapid generation times suited for secondary school laboratory classes, that can (4) foster links between school-based data and discoveries with a global community of professional researchers. These courses benefitted from accompanying web-based instruction (see below) and could be readily adapted to suit shorter or longer instructional timeframes. For example, after generating, then improving donor fly characteristics (**Fig 1**), and streamlining curricula (**Fig 6**), we have updated our course at two Stan-X partner schools. These modifications are perhaps better-matched to shorter instructional timeframes like summer terms, or the inclusion of fruit fly experiments as a part of an existing advanced biology class. Although we focused on frequently-used GAL4 lines in this study, university-based research laboratories could also nominate their own GAL4 lines for students to convert, thus fostering direct communication, and a feeling of ‘ownership’ and purpose in student collaborators.

To train instructors with little to no experience with *Drosophila* or CRISPR, we developed a week-long, intensive teacher training academy, called *Discover Now*. This approach of ‘teaching the teachers’ has fostered the autonomy of Stan-X instructors and their schools (Kim et al 2023; Chang et al 2022; Wendler et al 2022). Currently, partnering teachers from four additional schools are training to adopt HACKy-based experiments and instruction (S.P., N.L., unpubl. results). To provide practical guides for prospective research scientists and instructors interested in adopting this curriculum in their laboratory classes, the course manual is posted on the Stan-X website (https://www.stan-x.org/publications) and is periodically updated. In summary, we developed experiment-based courses to provide genuine science experiences to secondary school students while generating useful tools for the community of science. This experiential instruction has introduced the wonder, anxiety, and joy of scientific discovery to secondary school students, and informed their choices to pursue additional science training.

---

**Figure S1.**
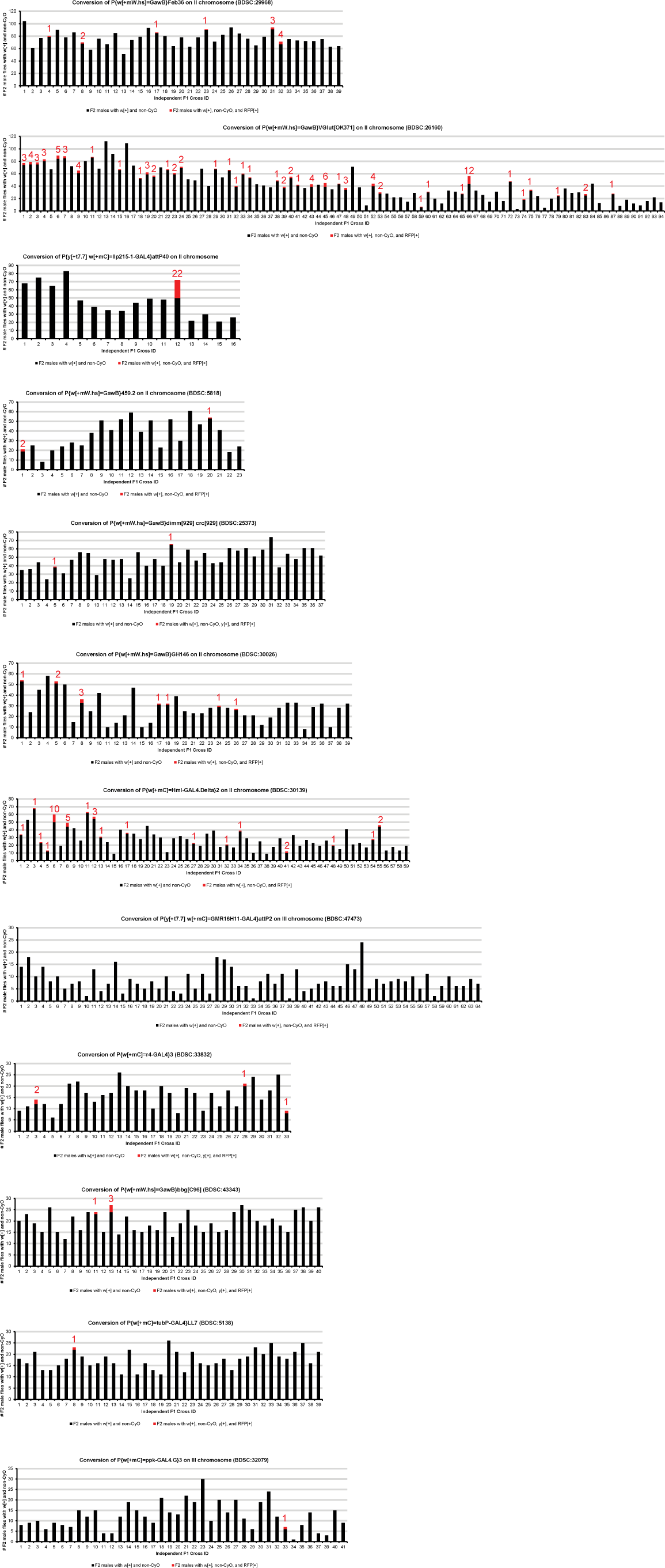
Frequencies of conversion events from independent male germlines. Phenotypic counting of F2 male progeny was plotted for each mating pair (Independent F1 Cross ID on X-axis). Black bars represent counts of F2 males with mini-white and non-curly phenotypes while red bars indicate F2 males with mini-white, non-curly, and RFP^+^, with frequencies written in red. For the second chromosome-linked GAL4 lines, each mating pair of a single F1 male and two *y*^1^ *w*^1118^ virgin females produced about 30 F2 males with desired phenotypes in a vial; the progeny size can be increased to 60 if F1 mating pairs are flipped once to a new vial after six days of initial mating. For X or third chromosome-linked GAL4 lines, these numbers decrease by a factor of one-half due to the independent segregation of donor and target chromosomes.

**Figure S2.**
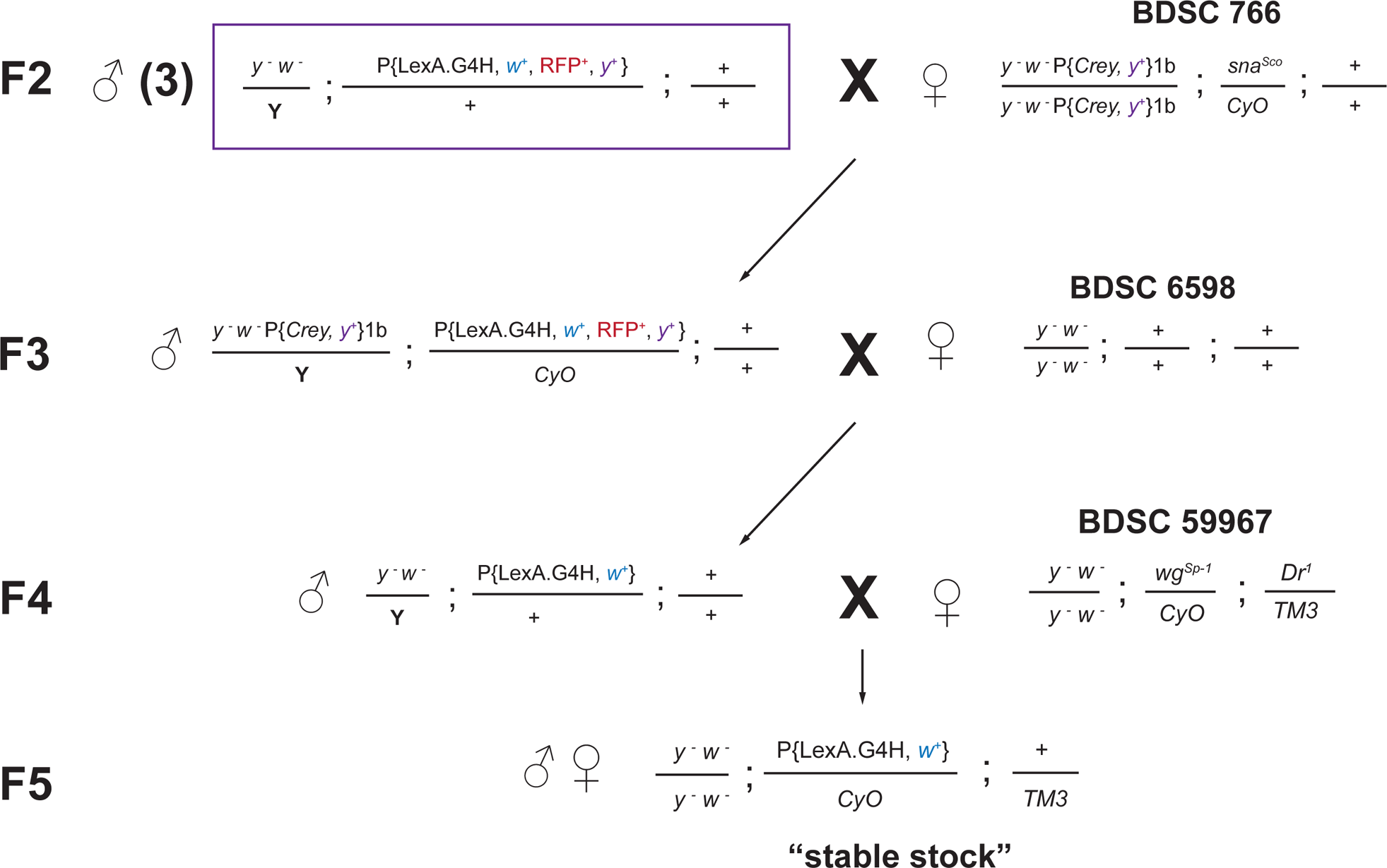
Mating scheme for removing loxP-flanked transgene cassette and establishing stable LexA.G4H lines. A single F2 male carrying the converted LexA.G4H transgene was mated to Cre-expressing virgin females (BDSC 766). A single F3 male with two transgenes was mated with virgin females of *y*^1^ *w*^1118^ (BDSC 6598). All F4 males carrying the mini-white transgene were without RFP and yellow transgene expression, but a single F4 male with the mini-white transgene was selected to mate with virgin females carrying balancer chromosomes (e.g. BDSC 59967) to isolate the chromosome with LexA.G4H transgene without RFP and yellow transgenes. In the F5 generation, the chromosome carrying LexA.G4H was balanced to establish a “stable stock” in the *y*^1^ *w*^1118^ genetic background.

**Movie S1. Live imaging of early pupa GFP expression in circulating hemocytes driven by either Hml-GAL4 (left) or Hml-LexA.G4H (right).** https://youtu.be/Bk EaKTiVE

## DATA AVAILABILITY STATEMENT

Strains and plasmids are available upon request. The course teaching materials and syllabuses are posted on the Stan-X website (https://www.stan-x.org/publications) and periodically updated. The authors affirm that all data necessary for confirming the conclusions of the article are present within the article, figures, and tables.

## ACKNOWLEDGMENTS

We thank the Bloomington Drosophila Stock Center (NIH P40OD018537) for fly stocks and FlyBase (NHGRI P41HG000739) for updated information. We thank past and current members of the Kim lab for helpful discussions and welcoming secondary school students. We thank S. Murray, I. Saxe, G. Hannon, M. Rupert, R. McGuire (Lawrenceville School), A. Hobbie and W. Rawson (Phillips Exeter Academy) for their advice, support, and encouragement. We are grateful to Philip Weissman (Micro-Optics Precision Instruments, NY) and Ken Fry (Genesee Scientific, CA) for generous support of equipment procurement for this project. We thank Glenn and Debbie Hutchins for supporting opportunities for students to engage in innovative science research at the Lawrenceville School.

## FUNDING

Work at Phillips Exeter Academy was supported by the John and Eileen Hessel Fund for Innovation in Science Education. The Hutchins Scholar Program at Lawrenceville School was supported by the Hutchins Family Foundation. Work in the Kim group was supported by NIH awards (R01 DK107507; R01 DK108817; U01 DK123743; P30 DK116074 to S.K.K.), the H.L. Snyder Foundation, the Elser Trust, gifts from Mr. Richard Hook and two anonymous donors, and the Stanford Diabetes Research Center.

## COMPETING INTERESTS

The authors declare no competing interests.

## AUTHOR CONTRIBUTIONS

AER, EF, TC, and NL were course instructors. AR and WP were teaching assistants. EG, JH, CS, MT, JW, AY, ESK, NKAA, PC, ACKL, MEL, JL, and KP were students. EW and PHC were undergraduate research assistants who generated final images. LK, SP, and SKK designed and managed the project. PHC, LK, SP, and SKK analyzed the data and wrote the manuscript. All authors read and approved the final manuscript.

